# Evening blue light exposure during adolescence induces avoidance behaviors and rewires medial amygdala circuit

**DOI:** 10.1101/2025.01.31.635947

**Authors:** Pablo Bonilla, David McBride, Alexandria Shanks, Janhvi Kartik, Ashita Bhan, Alessandra Porcu

## Abstract

Adolescents are being increasingly exposed to artificial blue light from electronic devices, raising concerns about its effects on brain development and mental health. The medial amygdala (MeA), a brain region critical for emotional regulation, is light-sensitive, yet how evening blue light during puberty influences its circuitry and behavior remains unknown.

Using a light cycle disruption paradigm, we found that adolescent mice exposed to evening blue light displayed increased avoidance behaviors compared to those exposed to darkness or reduced blue light conditions. Single-nuclei RNA sequencing revealed altered cell-type composition and disrupted synaptic communication pathways in the MeA. *In vivo* calcium imaging showed increased activity in MeA somatostatin neurons during avoidance behaviors, while chemogenetic inhibition of these neurons reduced these behaviors.

Our findings identify the MeA as a key integrator of emotional responses to environmental blue light, suggesting evening blue light exposure during puberty as a potential risk factor for affective disorders.

**Teaser:** Evening blue light exposure during adolescence alters medial amygdala functions, leading to anxiety-like behaviors.

## Introduction

Light is crucial for most species, and for eons, organisms have evolved to align their physiology and behaviors with the natural environmental light-dark rhythms, as human life has been historically synchronized to these predictable daily cycles. However, currently more than 80% of the world and more than 99% of the US and European populations live under light-polluted skies (*1*). Additionally, indoor light exposure has significantly increased in recent decades due to the widespread use of electronic devices and LED lighting, both of which emit blue light, disrupt natural nighttime darkness, and pose potential risks to human health (*2–5*).

Adolescents experience greater exposure to chronic light cycle disruption than any previous generation, as the use of electronic devices late into the night leads to a night owl pattern, with late bedtimes and early wake times to attend school (*6–9*). Recent studies have linked altered light environments to increased anxiety and mood disorders among U.S. adolescents (*10*, *11*). Pre-clinical studies have indicated that mice exposed to aberrant lighting conditions before adolescence develop avoidance behaviors in adulthood (*12*, *13*), suggesting that early life lighting environments play an important role in affective development. Emerging evidence suggests that altered light environments have consequences on brain development, potentially contributing to vulnerability to affective disorders (*14*, *15*). In our previous study, we developed a light paradigm for chronic light cycles disruption with evening blue light and implemented it in adolescent mice (*16*). This light paradigm specifically alters the diurnal activity of the medial amygdala (MeA), a key region involved in regulating emotional responses, such as innate avoidance and approach behaviors (*17*, *18*). Notably, the MeA is one of several brain regions receiving direct projections from intrinsically photosensitive retinal ganglion cells (ipRGCs) (*19*, *20*), which are responsible for non-image-forming light detection and are particularly sensitive to blue wavelengths (*21*, *22*). However, the impact of different wavelengths of light on the function of brain regions regulating emotional response, such as the amygdala (*23*), during adolescence when these regions are still maturing is not well understood.

Human studies have demonstrated that the spectral quality of light influences emotional responses in the amygdala (*24*, *25*). Specifically, blue light (compared to green) heightened responses to emotional stimuli and strengthened the functional connectivity between the voice area, amygdala, and hypothalamus. These findings indicate that the spectral composition of ambient light affects the processing of emotional stimuli, with blue light being particularly effective in engaging a network that integrates affective and ambient light information. Therefore, unveiling the neurobiological effects of different wavelengths of light during adolescence is of utmost significance for understanding the potential consequences on mental health.

Here, we show that adolescent mice exposed to light cycle disruption with evening blue light exhibit increased avoidance behaviors compared to mice exposed to the same protocol with reduced blue light or a control condition. Single-nuclei RNA sequencing revealed that exposure to evening blue light altered MeA neuronal circuits, including changes in neuronal composition and cell-cell communication within this area. Analysis of neuronal clusters showed alterations in the somatostatin (SST) neuronal population, including changes in gene expression that regulate neuronal activity. *In vivo* calcium recordings demonstrated that SST neurons exhibited increased neuronal activity associated with heightened avoidance behaviors. Selective chemogenetic inhibition of MeA SST neurons following evening blue light exposure was sufficient to reduce avoidance behaviors in adolescent mice.

Our findings uncover key molecular, cellular, and behavioral changes associated with evening blue light exposure during adolescence. Given that the MeA is conserved across species, our study may provide valuable insights into the fundamental biological processes disrupted by alterations in light cycles. Furthermore, we establish the MeA as a critical hub integrating environmental light and emotional responses, advancing our understanding of how light exposure during sensitive developmental periods may impact emotional regulation.

## Results

### Light cycle disruption with evening blue light but not reduced blue light exposure during adolescence increased innate avoidance behaviors in mice

Our previous study demonstrated that exposure to LCD during adolescence increased avoidance behaviors in mice (*16*). Studies in humans and rodents showed that the effects of light on emotional processing are wavelength-dependent (*25*, *26*). Therefore, we evaluated whether exposure to LCD with reduced blue light exposure (LCD-RBL) during adolescence alters avoidance responses in mice. To this aim adolescent mice (post-natal day 30) were exposed to control conditions (12 hours light:12 hours dark), a light cycle disruption paradigm with 12 hours of light followed by 5 hours of darkness and 7 hours of blue light (LCD-BL), or 7 hours of reduced blue light condition (LCD-RBL) for 4 weeks (Fig. 1A, Suppl. Fig. 1A-B). Mice (post-natal day 60) were then tested in the open field (OF) and elevated plus maze (EPM). In the OF test (Fig. 1B-D), adolescent mice exposed to LCD-BL showed decreased time spent in the center of the arena (inner zone) (Fig. 1B) and increased time spent in the perimeter (thigmotaxis) (Fig. 1C), while mice exposed to LCD-RBL spent significantly more time exploring the center of the arena and less time in the perimeter compared to the LCD-BL. In the EPM (Fig. 1E-G), adolescent mice exposed to LCD-BL showed reduced time spent in the open arms (Fig. 1E) and increased time spent in the closed arms of the maze compared to the control group (Fig. 1F). Remarkably, adolescent mice exposed to LCD-RBL spent significantly more time in the open arms (Fig. 1E) and less time in the closed arms (Fig. 1F) compared to the LCD-BL, showing the same exploration levels as the control group.

**Figure 1.**
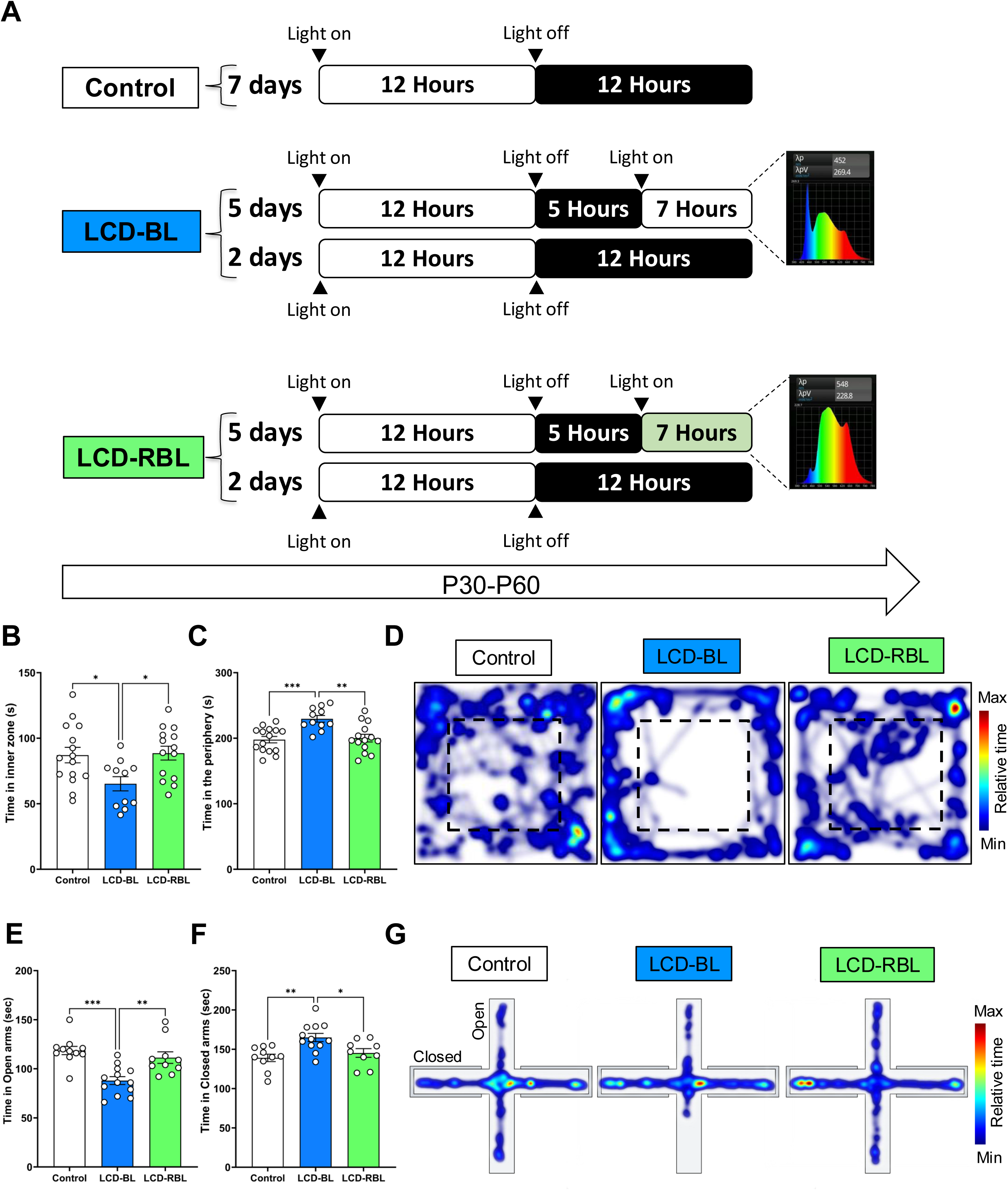
LCD-BL induces avoidance behavior in adolescent mice in the open field and the elevated plus maze. (**A)** Schematic representation of the light paradigms (light cycle disruption with evening blue light:LCD-BL; light cycle disruption with reduced evening blue light: LCD-RBL;. (**B)** Bar graph of time spent in the inner zone and (**C)** in the periphery of the open field arena. (**D)** Representative mouse position heatmaps in the open field. (**E)** Bar graph of time spent in the open arms and (**F)** in the closed arms of the elevated plus maze. (**G)** Representative mouse position heatmaps in the elevated plus maze. Individual data points represent independent mice, data are shown as mean ± SEM. One-way ANOVA Tukey multiple comparison post-test: \**p* < 0.01, \*\**p* < 0.01, \*\*\**p* < 0.001.

We also evaluated the effect of LCD-RBL exposure on the circadian locomotor activity rhythms recording voluntary wheel running activity for 3 weeks as shown in Suppl. Fig. 2A. Wheel running analysis revealed that mice exposed to LCD-RBL did not show differences in the total activity during the 7 hours of reduced blue light exposure (from ZT17 to ZT24) on the weekdays versus the 7 hours of dark phase on the weekends (Suppl. Fig.1B). We also observed similar activity onsets (Suppl. Fig. 2C) and total activity within the weekdays and weekends (Suppl. Fig. 1D). Additionally, adolescent mice exposed to LCD-BL and LCD-RBL showed no changes in activity in a novel environment compared to control mice (Suppl. Fig.1E). To test whether adolescence represents a critical window for change in light environment, we exposed adult mice (P70) to LCD-BL or Control conditions for 4 weeks and subsequently tested avoidance behavior in the EPM and OF. Adult mice exposed to LCD-BL did not exhibit significant differences in the inner zone of the open field test compared to control mice (Suppl. Fig. 3A) and in the time spent in the open arms of the elevated plus maze (Suppl. Fig. 3B). Altogether, our findings demonstrate that the increase in avoidance behavior observed after LCD exposure is both wavelength-dependent and specific to the adolescent developmental stage.

### Light cycle disruption with evening blue light rewires the MeA of adolescent mice

To study the impact of LCD with different light wavelengths on cellular states associated with avoidance behaviors, we performed snRNA-seq on MeA tissues from adolescent mice following 4 weeks of LCD-BL, LCD-RDL or control conditions (Fig.2A). We acquired a total of 13,764 single-nuclei transcriptomes mapped to a total of 21,136 unique genes from acutely microdissected and dissociated MeA cells. Based on the expression of previously known common cell-type markers, we identified six main cell types in the MeA: astrocytes, microglia, neuroblasts, neurons, oligodendrocytes and oligodendrocyte precursor cells (OPCs) (Fig. 2B and Suppl. Fig. 4). To assess the effects of evening blue light exposure on the different clusters, we conducted frequency analysis across these cell types (Fig. 2D). Frequency analysis of identified cell clusters revealed differences in the distribution of specific cell types across light conditions. Notably, astrocytes appeared reduced under LCD-BL compared to both controls and LCD-RBL conditions, while neurons appeared to increase, suggesting potential alterations in cellular composition associated with light exposure (Fig. 2D). Although these observations are based on a single biological replicate, they provide preliminary insights into how distinct light wavelengths may influence cellular states within the MeA. We then used immunofluorescence to validate these effects by quantifying the mature neuronal marker NeuN and glial fibrillary acidic protein (GFAP) in the MeA under the three light conditions (Supp. Fig. 5A-C). Interestingly, we observed no difference in the number of neurons; however, exposure to LCD-BL resulted in a decrease in the number of GFAP+ cells (Supp. Fig. 5B-D), corroborating observations in the snRNA-seq data. Given that astrocytes are crucial for cellular communication and the maintenance of network connectivity, we investigated whether the communication network in the MeA is altered following exposure to LCD-BL. To this end, we first classified the neuronal cluster into excitatory and inhibitory neurons (Suppl. Fig. 6) and then employed the CellChat v2 toolkit (*27*), to analyze cellular communication networks based on the expression levels of receptors and ligands across cell types. We found that LCD-BL exposure downregulates communication between excitatory neurons, inhibitory-excitatory neurons, and astrocytes (Fig. 2E). Additionally, exposure to LCD-RBL similarly downregulates excitatory neuron and inhibitory-excitatory neuron communication and astrocyte communication, while increasing microglial communication with OPCs, oligodendrocytes, and inhibitory neurons (Fig. 2E). These data suggest that exposure to LCD with different light wavelengths, particularly blue light, alters cellular composition and disrupts communication networks within the MeA, potentially influencing the neural mechanisms underlying avoidance behaviors.

**Figure 2.**
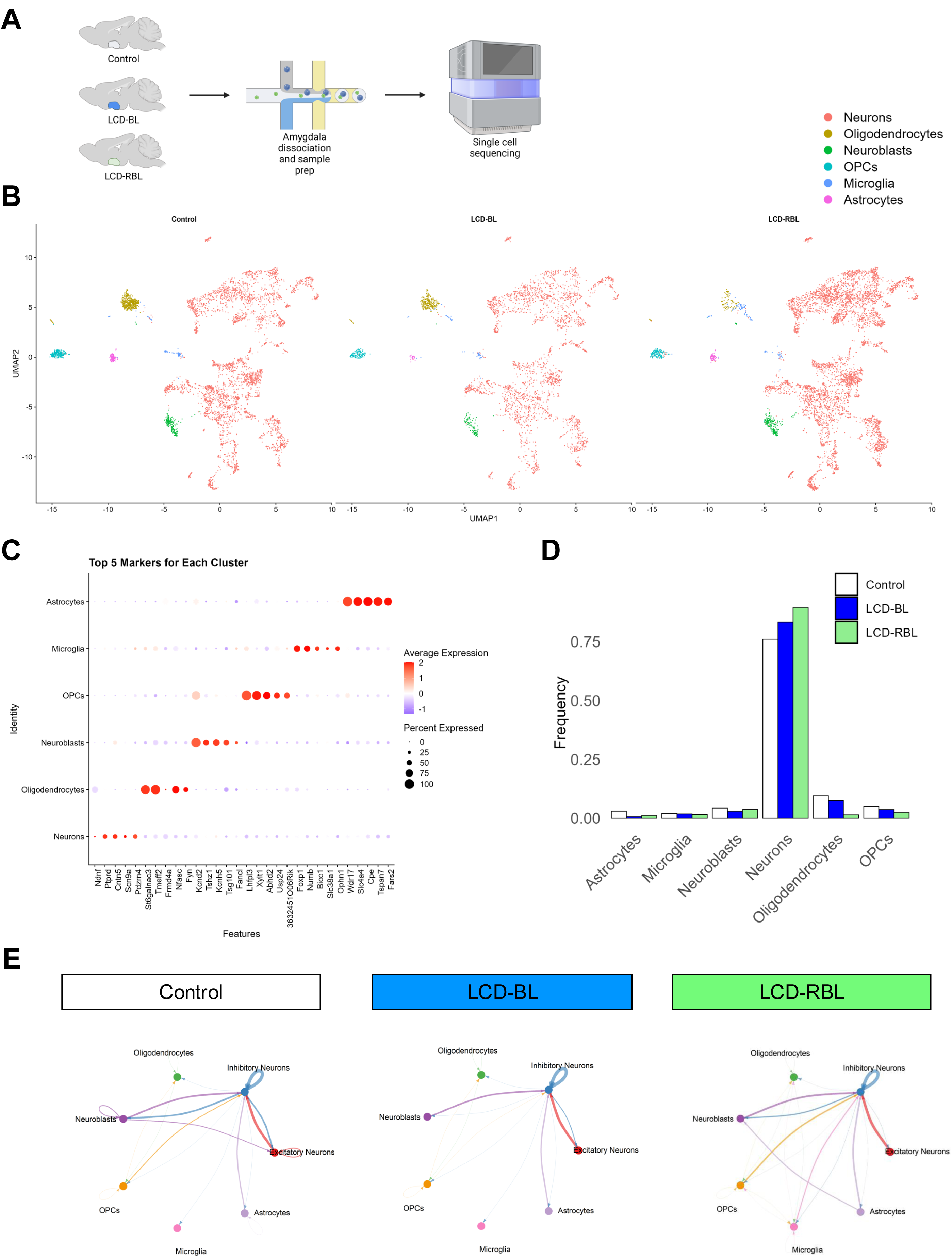
LCD-BL rewires the MeA of adolescent mice. **(A)** Schematic showing representative sagittal sections containing the dissected MeA region in the three experimental conditions, and single-nuclei isolation and sequencing using 10X Genomics. (**B)** UMAP of 13,764 number of MeA cells. Individual dots correspond to single cells colored by cluster identity according to major cell types identified in the MeA in Control, LCD-BL and LCD-RBL. (OPCs: Oligodendrocyte Progenitor Cells). (**C)** Dot plot of top 5 DEGs markers on the identified 6 cell clusters. **(D)** Bar graph showing the frequency of each cell type in Control, LCD-BL and LCD-RBL. **(E)** Visualizations of the inferred communication networks among multiple cell types of MeA. Circle plot is used to visualize the intercellular communication networks aggregated by method “weight”. In the circle plot, the node size indicates the strength of the outgoing signal from the cell, and the text labels are labeled by different colors to indicate their cell classes (i.e., Excitatory neurons, inhibitory neurons and non-neuronal); the communication strength for each link is indicated by the width.

### Neuron-specific clustering reveals altered sub-types and neuronal communication following exposure to light cycle disruption with evening blue light

To investigate neuroadaptations in the MeA following LCD-BL exposure, we identified four neuronal subtypes— two glutamatergic (*Slc17a6* and *Slc17a7*), one GABAergic (*Gad2*), and SST/Calbindin (*Sst/Calb1*) subtypes—based on the expression of established cell marker genes (Fig. 3A-B). Key markers of these subsets included well-known subpopulation genes that have been validated in prior reports (*28*). Frequency analysis revealed that LCD-BL exposure leads to a slight increase in SST/Calbindin neurons, accompanied by a corresponding decrease in glutamatergic (*Slc17a6*) neurons compared to LCD-RBL and control conditions (Fig. 3C-D). Notably, both LCD-BL and LCD-RBL exposures were associated with a reduction in *Gad2* neurons and an increase in glutamatergic (*Slc17a7*) neurons relative to the control condition (Fig. 3C-D). We then employed NeuronChaT (*29*) to infer intercellular communication among the identified neuronal subtypes. This analysis revealed that LCD-BL exposure moderately upregulated *Slc17a7*-*Slc17a6* and *Slc17a6*-SST/Calbindin signaling compared to control conditions (Fig. 3E). Interestingly, LCD-RBL exposure appeared to downregulate *Slc17a7*-*Slc17a6* communication while upregulating *Slc17a6*-*Gad2* signaling, suggesting an enhanced GABAergic function within the MeA (Fig. 3E).

**Figure 3.**
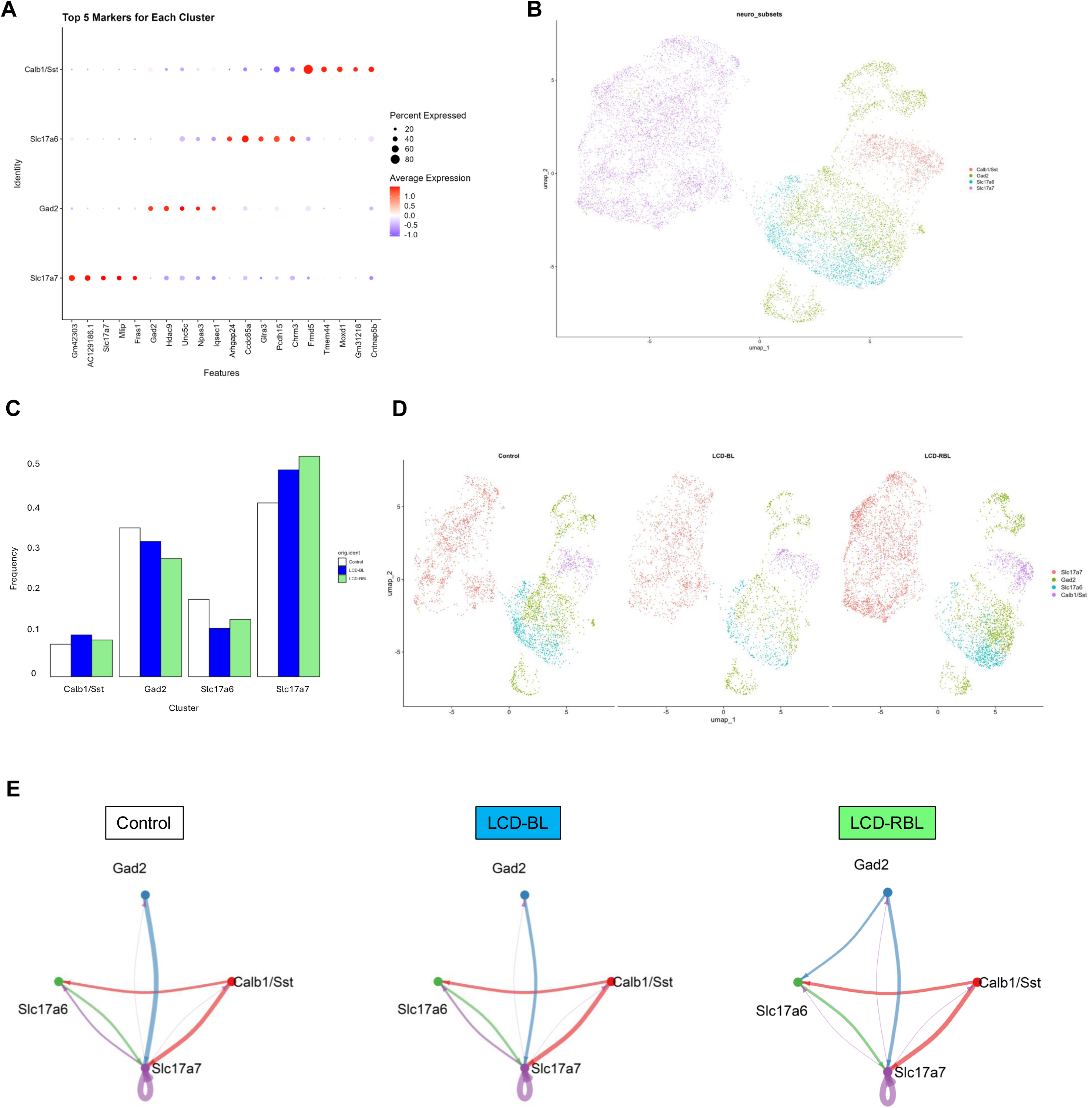
LCD-BL alters neuronal subtypes distribution and neuron-neuron communication in the MeA of adolescent mice. **(A)** Dot plot representative subtype expression level markers across the 4 neuronal subclusters Calb/SST, Slc17a6, Gad2 and Slc17a7. (**B)** UMAP representation of 11,476 of neurons colored by neuronal subcluster identity. (**C)** Bar graph shows the frequency of Calb/SST, Slc17a6, Gad2 and Slc17a7 neuronal subclusters in control, LCD-BL and LCD-RBL conditions. **(D)** UMAP representation of 11,476 neurons colored by neuronal subcluster identity split by treatments from control, LCD-BL and LCD-RBL conditions. (**E)** Visualizations of the inferred communication networks among multiple neuronal subtypes in the MeA. Circle plot is used to visualize the intercellular communication networks aggregated by method “weight”. In the circle plot, the node size indicates the strength of the outgoing signal from the cell, and the text labels are labeled by different colors to indicate their cell classes (i.e., Calb/SST, Slc17a6, Gad2 and Slc17a7); the communication strength for each link is indicated by the width.

**Figure 4.**
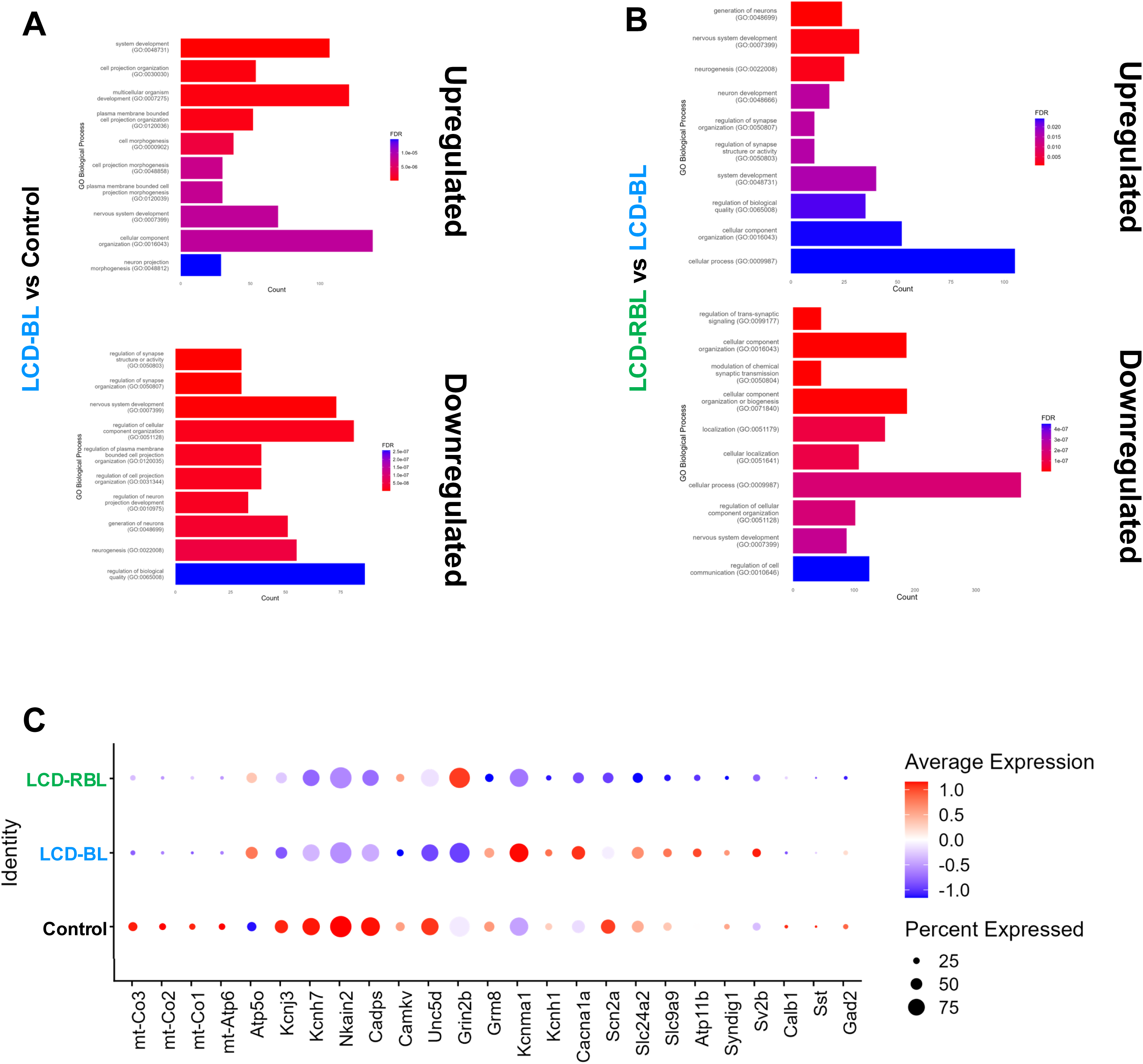
LCD-BL influences the expression of genes regulating synaptic transmission and neuronal activity. **(A)** Top gene ontology terms (FDR < 0.15) upregulated and downregulated in the neuronal population in LCD-BL vs control. (**B)** Top gene ontology terms upregulated and downregulated in the neuronal population in LCD-RBL vs LCD-BL. (**C)** Dot plot showing the distribution of expression level of selected genes involved in synaptic transmission and neuronal activity. Dot size indicates percent expressed while color indicates average expression.

### Gene ontology and differential gene expression uncover molecular changes in neuronal subtypes following exposure to light cycle disruption with evening blue light

To further explore the molecular changes underlying the observed neuroadaptations, we analyzed the effect of LCD-BL exposure on gene ontology (GO) enrichment and differential gene expression within neuronal subtypes. This approach allowed us to identify key pathways and biological processes impacted by light cycle disruption with evening blue light. Top enriched GO term analysis revealed distinct biological processes altered by LCD-BL exposure compared to the control group. Upregulated processes included nervous system development and cell projection morphogenesis, while down regulated processes were enriched in genes associated with synapse structure, organization, and activity, as well as neurogenesis and cell projection morphogenesis (Fig. 3A). Further molecular function analysis identified the top four upregulated processes as GTPase regulator activity, nucleoside-triphosphatase regulator activity, guanyl-nucleotide exchange factor activity, and small GTPase binding (Suppl. Fig. 7A). The top four down regulated processes included phosphatase binding, cell adhesion binding, transcription coregulator activity, and protein phosphatase binding (Suppl. Fig. 7B). In contrast, the LCD-RBL group showed upregulated biological processes related to neurogenesis, development, and regulation of synapse structure and activity compared with LCD-BL (Fig. 3B). Downregulated processes in the LCD-RBL group included those involved in synaptic transmission, cellular component organization and localization, and nervous system development (Fig. 3B). The top four upregulated molecular functions were protein serine/threonine kinase activity, GTPase regulator activity, nucleotide-triphosphatase regulator activity, and guanyl-nucleotide exchange factor activity (Suppl.Fig. 7C). The top four down regulated processes included phosphatase binding, protein serine/threonine kinase activity, GTPase binding, and transmembrane transporter binding (Suppl. Fig. 7D). Overall, these findings suggest that LCD-BL exposure disrupts synaptic transmission while LCD-RBL exposure preserves synaptic functionality and organizational integrity.

Differential gene expression analysis revealed that LCD-BL exposure altered the expression of genes involved in synaptic transmission and neuronal activity, including mitochondrial genes (*mt-Co1, mt-Co2, mt-Co3*), potassium channels (*Kcnj3, Kcnh1, Kcnh7, Kcnma1*), sodium channels (*Nkain2*), calcium channels (*Cacna1a*), and glutamate receptors (*Grin2b, Grm8*) (Fig. 3C). Notably, *Kcnj3*, which encodes a G protein-coupled inwardly rectifying potassium channel (GIRK), was significantly downregulated in LCD-BL mice compared to LCD-RBL and control groups. Given that different neuronal subtypes in the MeA regulate avoidance behavior (*18*) and that our earlier findings showed that MeA^SST+^ neurons are light-responsive (*16*), we next evaluated which neuronal population is specifically affected by LCD-BL exposure. Among the identified genes, we focused on *Kcnj3* due to its critical role in regulating neuronal activity and behavioral responses, including avoidance behaviors (*30*)*. In situ* RNAscope (RNA-ISH) analysis revealed that *Kcnj3* was significantly downregulated in the MeA^SST+^ of LCD-BL mice compared to control and LCD-RBL mice (Suppl. Fig. 8A-B). Furthermore, exposure to LCD-RBL increased expression of Kcnj3 in MeA^GABA+^ neurons compared to the LCD-BL (Suppl. Fig. 8C-D). No differences were found on the MeA^GLUT+^ neurons (Suppl. Fig. 8E-F). These findings suggest that the downregulation of *Kcnj3* in LCD-BL mice specifically impacts MeA^SST+^ neuronal activity, potentially contributing to changes in avoidance behaviors.

### Evening blue light-induced avoidance behavior is associated with increased MeA^SST+^ neuronal activity

We then hypothesized that avoidance behaviors are associated with increased MeA^SST+^ neuronal activity in adolescent mice after LCD-BL exposure. To this aim, following 4 weeks of control, LCD-BL or LCD-RBL exposure adolescent mice were sacrificed 45 minutes after EPM and OF tests (Fig. 5A). Brains were then processed for neuronal activity marker (c-FOS) analysis by immunofluorescence and RNA-ISH. We found that adolescent mice exposed to LCD-BL showed increased c-FOS expression in MeA^SST+^ neurons during EPM and OF (Fig. 5B-C) compared to control mice. Remarkably, adolescent mice exposed to LCD-RBL showed a significant reduction in c-FOS expression in MeA^SST+^ neurons compared to the LCD-BL group. Interestingly, neither MeA^GABA+^ (Fig. 5D-E) nor MeA^GLUT+^ (Fig. 5F-G) neurons exhibit significant differences in neuronal activity. These results imply that selective subtypes of MeA neurons are affected by altered light environments during adolescence.

**Figure 5.**
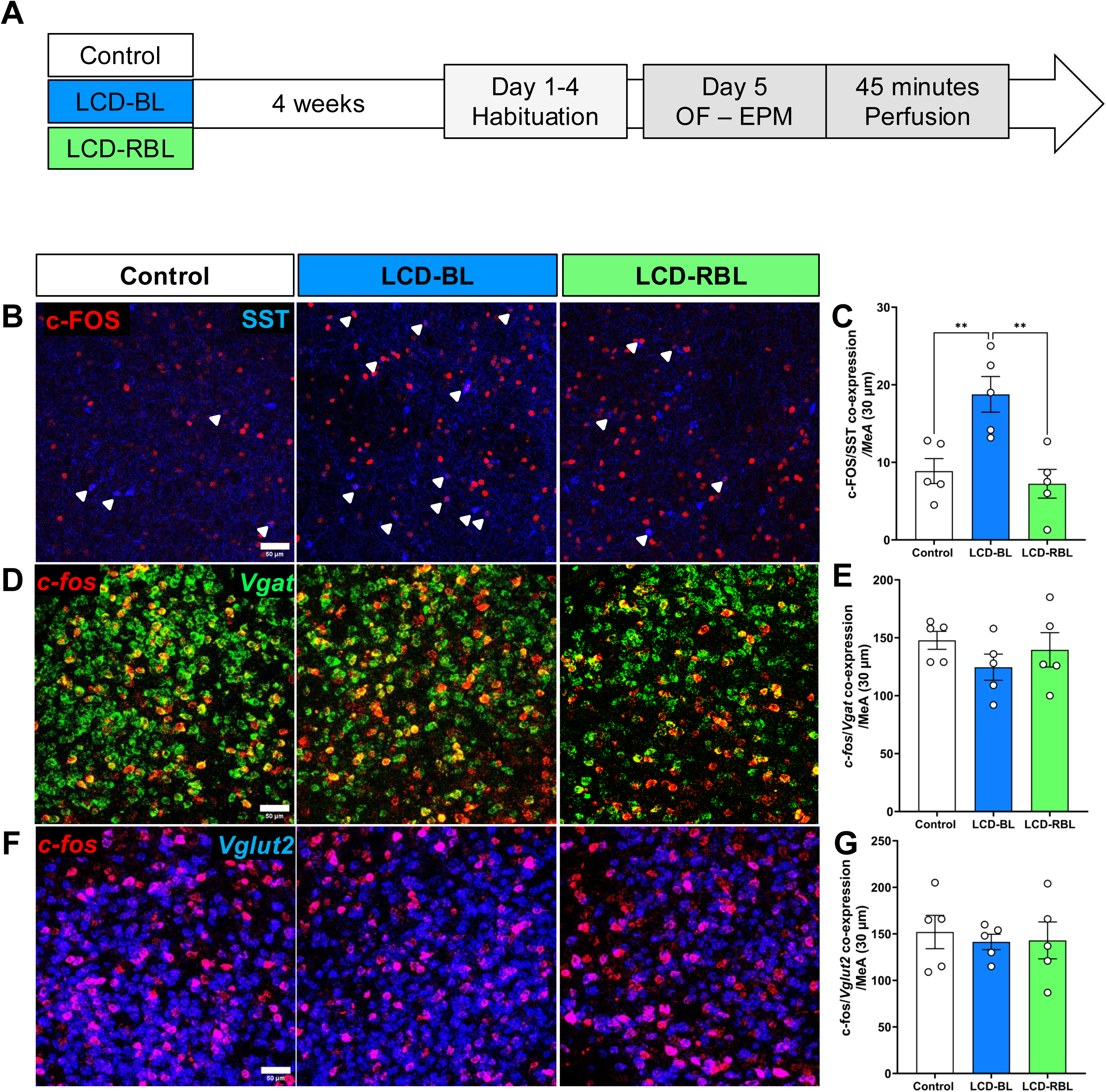
LCD-BL-induced avoidance behaviors are associated with increased MeA^SST+^ neuronal activity. **(A)** Schematic overview of the experimental design. (**B**) Representative confocal micrographs showing c-FOS (red) detected by immunofluorescence in SST neurons (SST, blue) and *c-fos* (red) mRNA expression detected by RNAscope (**D)** GABAergic neurons (*VGAT*, green) and (**F)** glutamatergic neurons (*VGLUT2*, blue) in the MeA (Scale bar 50 μm). Bar graph of c-FOS co-expression in SST+ neurons (**C**), GABAergic (**E**) glutamatergic neurons (**G**) in the MeA of adolescent mice exposed to control, LCD-BL and LCD-RBL conditions. Individual data points represent independent mice, data are shown as mean ± SEM. One-way ANOVA Tukey multiple comparison post-test: \*\**p* < 0.01.

To investigate the neuronal activity of MeA^SST+^ neurons *in vivo*, we used fiber photometry to record behavioral-evoked MeA^SST+^ activity in mice using a Ca^2+^ sensor (GCaMP8m) selectively expressed in SST neurons in the MeA during EPM and OF tests following 4 weeks of control, LCD-BL or LCD-RBL exposure (Fig. 6A-C). In the OF, adolescent mice exposed to LCD-BL showed increased Ca^2+^ activity in MeA^SST+^ neurons during the start of the exploration in the inner zone compared to control, as shown by GCaMP8m signal traces (Fig. 6E), area under the curve and mean value (Fig. 6F-G). In the EPM, adolescent mice exposed to LCD-BL showed increased Ca^2+^ activity in MeA^SST+^ neurons before and during the open arm exploration (Fig. 6I-K) compared to control. No significant differences were found in the maximum mean value, minimum mean value, or amplitude across conditions (Suppl. Fig. 9A-L). These results indicate that LCD-BL induced hyperactivity of MeA^SST+^ neurons which might drive avoidance responses in adolescent mice.

**Figure 6.**
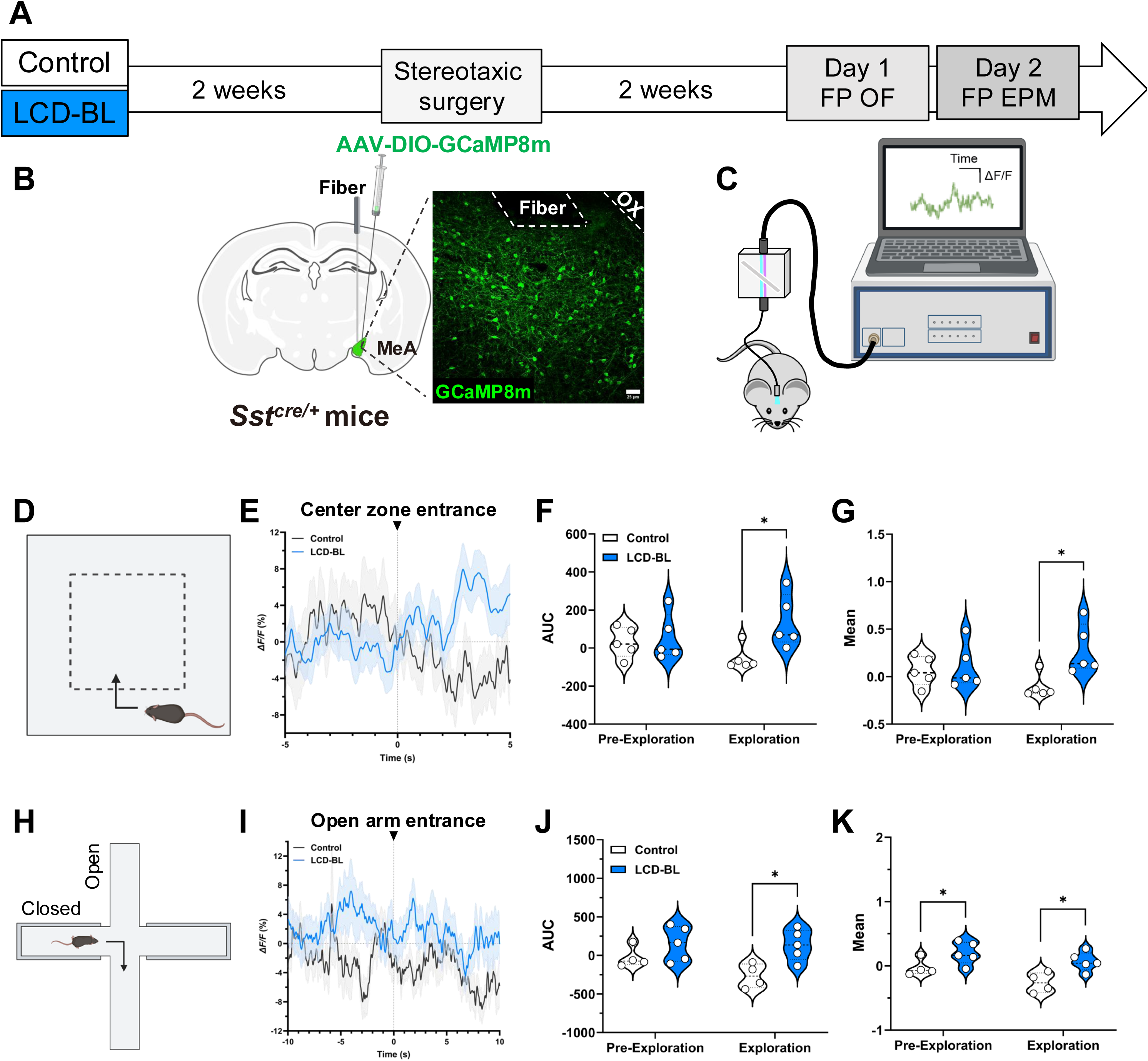
LCD-BL exposure increases MeA^SST+^ activity during explorations in the open field and elevated plus maze. **(A)** Schematic timeline showing the AAV intracranial injection for the fiber photometry recordings during Open Field (OF) and Elevated Plus Maze (EPM) tests. (**B)** Schematic of AAV injection and optic fiber implant in the MeA of SST-Cre mice, with confocal image showing GCaMP8m-expressing SST neurons (green) (Scale bar 25 μm). (**C)** Schematic of the fiber photometry system and recording. (**D)** Overview of exploratory events analyzed in the OF test. (**E)** GCaMP8m signal traces (% ΔF/F) during exploration, with event start indicated by a dotted line and arrowhead for event start. Violin plots show area under the curve (AUC) **(F)** and mean values (**G)** of GCaMP8m before (Pre-Exploration) and during the exploration (Exploration). **H)** Overview of exploratory events analyzed in the EPM test. (**I)** GCaMP8m signal traces (% ΔF/F) in MeA SST neurons during exploration, with event start indicated by a dotted line and arrowhead for event start. Violin plots show AUC (**J)** and mean values (**K)** of GCaMP8m before (Pre-Exploration) and during the exploration (Exploration). Individual data points represent independent mice, data are shown as mean ± SEM. Student’s t-test: *p<0.05.

### hM4Di-mediated inhibition of the MeA^SST+^ neurons suppresses avoidance behavior following evening-blue light exposure

To determine whether the enhanced MeA^SST+^ activity is causally related to the avoidance behavior observed in adolescent mice following 4-weeks of LCD-BL, we performed an acute chemogenetic inhibition of MeA^SST+^ neurons before the OF and EPM tests. To this aim, SST-Cre mice exposed to LCD-BL were stereotaxically bilaterally injected in the MeA using a Cre-dependent AAV encoding either an inhibitory DREADDs (AAV-Flex-hM4D-Gi) or a control virus (AAV-Flex-mCherry) and received an i.p injection with either JHU or saline 30 min prior to the behavioral tests (Fig.7A-B). In the OF test (Fig. 7C-E), GiDREADDs-JHU mice showed an increased time spent in the inner zone of the arena (Fig. 7C) and a significant reduction in the time spent in the periphery (Fig. 7D) compared to GiDREADDs-Saline or mCherry-JHU mice. Additionally, GiDREADDs-JHU mice showed decreased distance travelled compared to GiDREADDs-Saline mice (Suppl. Fig. 10A). In the EPM (Fig. 7F-H), GiDREADDs-JHU mice showed increased time spent in the open arms compared to GiDREADDs-Saline or mCherry-JHU (Fig. 7F) and reduced spent time in the closed arms compared to DREADDs-Saline (Fig. 7G). No significant differences were observed in the distance traveled (Suppl. Fig. 10B). Mice were then sacrificed 45 minutes after OF and EPM and brains were processed for neuronal activity marker (c-FOS) analysis by immunofluorescence. As expected, we found that GiDREADDs-JHU mice showed significantly decreased c-FOS expression in MeA^SST+^ neurons compared to GiDREADDs-Saline or mCherry-JHU mice following behavioral tests (Figure 7I-J). Additionally, GiDREADDs-JHU mice showed a decrease in total c-FOS expression (Suppl. Fig. 11A-B), suggesting that acute inhibition influences the MeA network. We conclude that the acute inhibition of the MeA^SST+^ neurons rescues the avoidance behavior induced by LCD-BL exposure, suggesting that MeA^SST+^ neurons are necessary for light-induced avoidance behaviors in adolescent mice.

**Figure 7.**
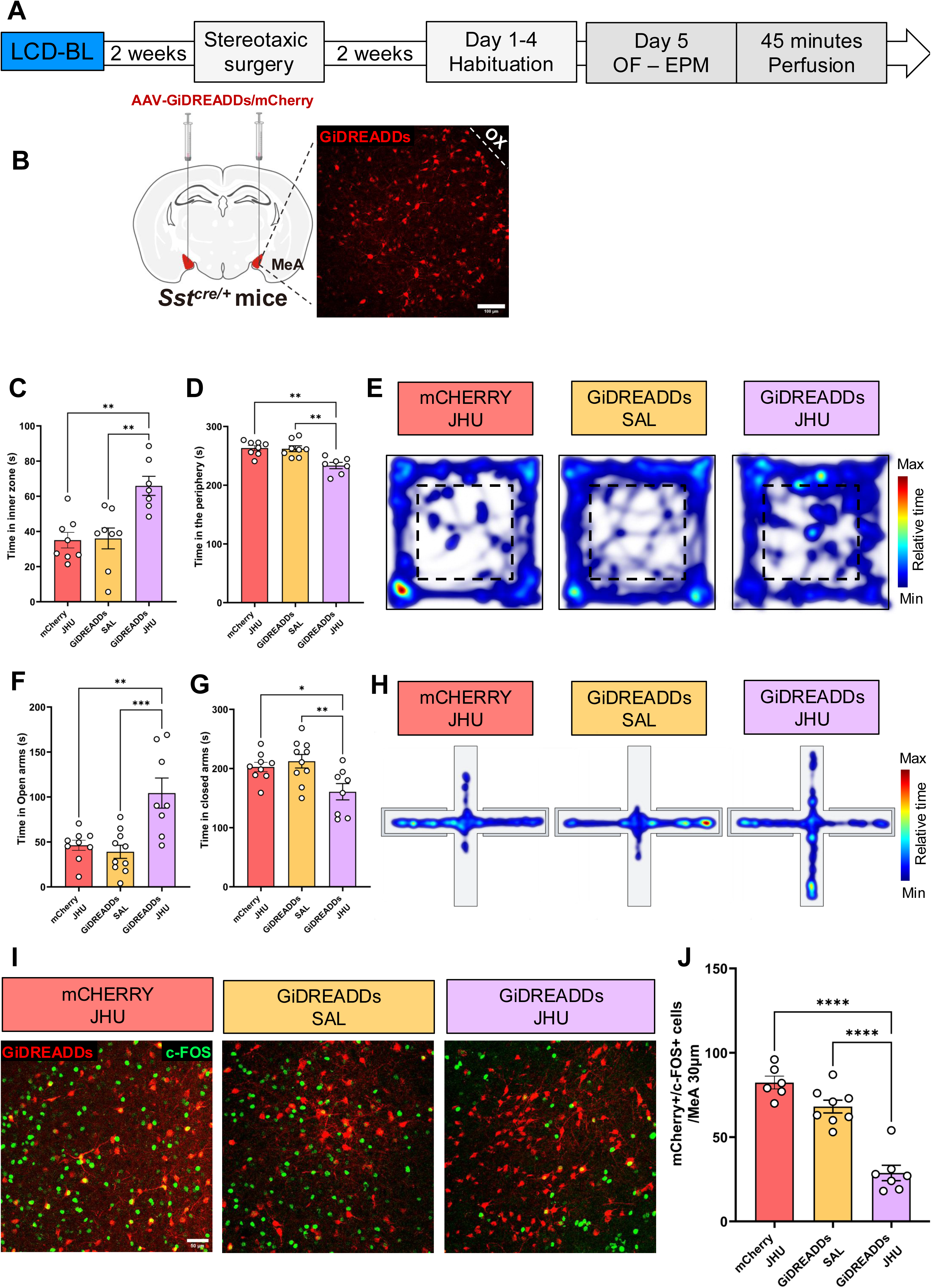
Acute inhibition of MeA^SST+^ neurons rescued LCD-BL-induced avoidance behavior in the open field and elevated plus maze. **(A)** Schematic timeline showing the DREADDs intracranial injection, habituation and behavior days in the Open Field (OF) and Elevated Plus Maze (EPM) and time of the perfusion. (**B)** Schematic of AAV bilateral injection in the MeA of SST-Cre mice, with confocal image showing GiDREADDs-expressing SST neurons (Red) (Scale bar 100 μm). (**C)** Bar graph of time spent in the inner zone and in the periphery (**D)** of the open field arena. (**E)** Representative heatmaps in the open field. Bar graphs of time spent in the open arms (**F**) and in the closed arms (**G)** of the elevated plus maze. (**H)** Representative heatmaps in the elevated plus maze. (**I)** Representative confocal micrographs showing c-FOS expression in MeA^SST+^ neurons expressing Gi DREADDs (Scale bar 50 μm). (**J)** Bar graph of c-FOS/mCherry co-expression in the MeA of adolescent mice after the open field and elevated plus maze test. Individual data points represent independent mice, data are shown as mean ± SEM. One-way ANOVA Tukey multiple comparison post-test: \**p* < 0.05, \*\**p* < 0.01. ns = Non significant.

## Discussion

We previously showed that exposure to light cycle disruption during adolescence altered emotional responses in mice without affecting the circadian pacemaker (*16*). In this study, we extend those findings by showing that light cycle disruption with specific wavelengths, particularly blue light, has profound effects on adolescent brain function and behavior. Specifically, we found that evening blue light exposure: (1) induces robust avoidance behaviors in adolescent mice, (2) rewires MeA circuits by altering cell-type composition and intercellular communication, and (3) increases MeA^SST+^ neuronal activity during avoidance responses. Notably, our chemogenetic inhibition experiments demonstrate that MeA^SST+^ neuronal activity is required to drive blue light-induced avoidance behavior. Together, these results implicate the MeA as a hub integrating emotional responses and environmental light and suggest that evening blue light exposure during puberty might represent a potential risk factor for abnormal amygdala development and affective disorders.

Importantly, the avoidance behaviors observed after four weeks of LCD-BL exposure were evident only when the light paradigm was applied during adolescence (P30–P60) and not during adulthood (P70–P100). This age-specific vulnerability is consistent with previous studies suggesting that adolescents are more sensitive to light exposure than adults (*31*). Furthermore, recent research has shown that irregular environmental lighting during adolescence is associated with increased anxiety disorders (*10*). These findings highlight adolescence as a critical developmental window during which environmental factors, such as aberrant light exposure, can significantly influence emotional and behavioral outcomes. In addition, a previous study revealed that mice exposed to dim light at night during the first three weeks of post-natal life exhibit increased anxiety as adults (*13*). Future studies should focus on the long-term effects of adolescent light exposure, investigating whether the behavioral and neural alterations persist into adulthood and contribute to the development of affective disorders later in life.

While previous research has examined how factors such as timing, luminance levels, and light exposure duration influence sleep-related responses in mice and alertness in humans (*32*, *33*), surprisingly few studies have investigated the impact of light wavelength on emotional responses. Notably, Vandewalle et al. found that blue light, compared to green light, enhanced responses to emotional stimuli and strengthened the functional connectivity between the voice area, amygdala, and hypothalamus (*25*). In line with these findings, our study shows that LCD with evening blue light exposure induces avoidance behaviors and alters MeA circuit function, compared to LCD with reduced blue light. These results suggest that wavelength-specific light exposure, particularly blue light, plays a critical role in modulating emotional processing and further underscore the importance of considering light wavelength in studies of light’s effects on brain function and behavior. Anatomical connectivity could easily support the differential responses to various light wavelengths in the MeA, with a preferential reaction to blue light likely due to direct inputs from ipRGCs (*19*). These direct ipRGC inputs to the MeA may facilitate the increased sensitivity to blue light observed in our study. However, other subnuclei of the amygdala, such as the central amygdala (CeA), have recently been shown to modulate anxiety-like behavior in response to acute high-intensity light exposure (*34*). This suggests that light exposure may engage different amygdala circuits depending on the intensity and wavelength of light, potentially leading to distinct emotional and behavioral outcomes. While we suggest that the MeA appears to play a central role in processing blue light exposure, the CeA and other subnuclei may contribute to the broader context of light-induced anxiety and emotional regulation. Future studies should explore how these distinct amygdala subnuclei interact under varying light conditions and whether they work in concert or independently to regulate emotional responses.

In contrast to adults, the effects of altered light environments during developmental stages particularly when brain regions that regulate emotional responses, such as the amygdala, are still maturing remain poorly understood. To address this gap, we implemented snRNA sequencing to provide, for the first time, insights into how different light wavelengths impact the maturation and function of MeA circuits. One striking finding from our study is the profound impact of LCD with blue light exposure on intercellular communication among MeA cell types and neuronal subpopulations. Specifically, communication between inhibitory and excitatory neurons in the MeA was significantly altered in response to blue light exposure. These changes in the balance of excitation (*Slc16a and Slc17a*) and inhibition (*Gad2*) within the MeA are particularly noteworthy, as inhibitory and excitatory subpopulations in this region play key roles in controlling opposing approach/avoidance responses (*17*, *18*). The disruption of this delicate balance could potentially contribute to altered emotional responses, such as increased avoidance behaviors observed following LCD exposure. In addition, LCD with blue light exposure affects both glial and neuronal subtype composition within the MeA. Specifically, we observed alterations in the proportions of astrocytes, which are essential for maintaining proper synaptic function, modulating neuronal activity, and ensuring circuit stability (*35*). These changes could impair the MeA’s ability to process and integrate emotional stimuli, further contributing to the increased avoidance behaviors. Given the critical role of astrocytes in supporting synaptic plasticity and neuronal communication (*36–38*), future studies should focus on the specific contributions of astrocytes in these processes.

Our differential gene ontology analysis revealed significant changes in biological processes following exposure to LCD with blue light paradigm. Upregulated processes included those related to nervous system development and cell projection morphogenesis, further indicating potential remodeling of neural circuits. In contrast, downregulated processes were enriched in genes associated with synapse structure, organization, and activity. These findings suggest that LCD with blue light exposure may impair key processes involved in maintaining synaptic stability and plasticity, while promoting developmental changes that could affect neural function. Indeed, differential gene expression analysis revealed that exposure to LCD with evening blue light paradigm altered the expression of several genes critical for synaptic transmission and neuronal activity, such as *Kcnj3*, which encodes GIRK (*39*). GIRK channels activation hyperpolarize neurons in response to activation of many different G protein-coupled receptors and thus control the excitability of neurons through GIRK-mediated self-inhibition, slow synaptic potentials, and volume transmission (*39*). Moreover, GIRK activity in the amygdala has an important role in behavioral responses, including avoidance behaviors (*30*). Further investigation into GIRK channel expression and its role in regulating neuronal activity within the MeA during adolescence could uncover new targets for therapeutic strategies aimed at addressing affective symptoms induced by chronic light cycle disruption.

Our cFOS quantification and fiber photometry analysis further supported this, revealing that exposure to LCD with blue light paradigm increased MeA^SST+^ neuronal activity during avoidance behaviors. The heightened activity of MeA^SST+^ neurons in response to LCD with blue light exposure suggests that this specific neuronal subtype may play a key role in mediating the behavioral outcomes observed in our model. To test whether increased MeA^SST+^ activity is necessary for light-induced avoidance behavior, we performed acute chemogenetic inhibition of MeA^SST+^ neurons. This inhibition effectively rescued the avoidance response observed in the LCD with evening blue light paradigm, underscoring the critical involvement of MeA^SST+^ neuronal activity in driving these behavior changes. These findings are consistent with previous studies demonstrating that SST-expressing neurons in the amygdala play a pivotal role in mediating avoidance behavior (*40–43*). Furthermore, enhanced activity of SST neurons has been shown to be both necessary for and sufficient to drive stress-induced avoidance behavior (*44*), highlighting the role of MeA^SST+^ neurons in the regulation of emotion-driven behaviors.

Of note, a previous study found increased MeA^SST+^ activity following exposure to a new, stressful environment (*45*), further suggesting that SST+ neurons in the MeA are critical for adapting to and processing environmental stressors. This increased activity in response to stress and our findings of elevated MeA^SST+^ neuronal activity following exposure to the LCD with evening blue light paradigm suggest that SST neurons in the MeA might act as a key regulator, capable of adjusting to altered light environments to modulate avoidance behavior.

The study presented here expands our understanding of the MeA as a central hub for integrating emotional responses to environmental stimuli. We propose that evening blue light exposure during a critical developmental window may disrupt normal MeA circuitry, potentially increasing the risk of affective disorders. Additionally, we demonstrate that selective neuronal populations in the MeA drive emotional responses to specific light wavelengths, suggesting that these stimuli activate unique neuronal circuits within this brain area. This research underscores the significance of understanding how environmental factors, such as artificial light exposure, can influence brain development and mental health during adolescence. Future investigations should explore whether these findings are applicable to human populations and evaluate potential interventions to mitigate the effects of blue light exposure on mental health in teens.

## Materials and Methods

### Mice

Female and male mice used in this study were on a C57BL/6 J genetic background, except for GAD67-GFP mice that were on CD1 background. The Gad1-tm1.1Tama (GAD67-GFP knock-in) mouse line was provided by Y. Yanagawa (Gunma University Graduate School of Medicine, Japan). Mice were heterozygous for insertion of the gene encoding GFP under the control of the GAD67 gene promoter. They were used to label the inhibitory GABAergic neurons by enhancing the GFP signal with an anti-GFP antibody. For fiber photometry experiments *Sst^tm2.1(cre)Zjh^*/J mice (JAX #013044) were used. Mice were bred and reared in an on-site animal facility at UofSC under controlled environmental conditions and maintained in standard group housing cages with *ad libitum* access to normal chow and water. Light intensity was approximately 600 lux at the top of the cage, and the ambient temperature was 23° ± 2°C. The time referred to as “lights on” is defined as ZT0. We minimized the number of animals used in each experiment (sample size based on power analysis) and the animals’ pain and distress. Mouse studies were conducted in accordance with regulations of the Institutional Animal Care and Use Committee at UofSC (Protocol #2615-101736-090222).

### Chronic light cycle disruption paradigm

Adolescent mice (post-natal day 30) were transferred into light-tight circadian cabinets (Actimetrics, IL, United States) and subjected to one of the following lighting conditions for a period of 4 weeks: (1) standard 12:12 h light–dark cycles (12 L:12D) (light at ∼600 lux) (Control group); (2) 19 h of light and 5 h of darkness for 5 days and 12 L:12D for 2 days per week (LCD-BL group); (3) 12 h of light, 7 h of reduced blue light and 5 h of darkness, for 5 days and 12 L:12D for 2 days per week (LCD-RBL group) (Figure 1). LED light (∼600 lux) is turned on during the night phase of the animals’ cycle, resulting in increased duration of light exposure for 5 days of a 7-day cycle. Light spectrums and chromaticity are shown in Supplementary Figure 1. After 4 weeks of either LCD-BL, LCD-RBL or control conditions, mice were placed in 12 L:12D for 5 days and tested for exploratory behaviors and then sacrificed. 6 cohorts of mice were used: one for avoidance behaviors tests, one for snRNA-sequencing, one for histological analysis, one to evaluate cFOS expression after behavior, one for fiber photometry experiments and another one for the chemogenetic inhibition of MeA^SST+^ neurons.

### Behavioral assays

All mice were handled daily for at least one week before behavioral assays to ensure acclimation. During this period, they were placed in the behavior room for 15–20 minutes between ZT8 and ZT10. All behaviors were recorded and analyzed using EthoVision XT (Noldus Information Technology, USA) to track the location of mice. For the elevated plus maze test, mice were placed in a plus shaped plastic maze (Noldus Information Technology, USA), consisting of two open and closed arms (35×5 cm) extending from a central platform elevated from the ground by 50 cm. Each mouse was initially placed in the center of the maze facing the open arm of the maze and away from the experimenter and the behavior was recorded for 5 min. For the open field test, mice were placed in an open field chamber (40×40×40 cm), where the center zone was defined as a square at the center (28×28 cm). Each mouse was placed at the center of the arena at the beginning of the session.

### Stereotaxic surgery for virus injection and cannula implants

Male and female 6 weeks old *Sst^tm2.1(cre)Zjh^*/J mice were deeply anesthetized with 5% isoflurane and placed in a robot stereotax stereotaxic apparatus (Neurostar, Model 900HD Motorized Small Animal Stereotaxic) while resting on a heat pad under 2.5-1.5% isoflurane anesthesia. Following craniotomy, 650 nl of AAV5.Syn.Flex.GCaMP8m.WPRE.SV40 (AddGene, 162378), were injected unilaterally into the left MeA (AP: −1.58; L: −1.89; DV: 5.47 mm) using a 1 mL syringe (65458–01, Hamilton Company, Reno, NV; Stoelting). To allow time for diffusion of the virus, the injection needle remained immobile for 10 min before removal. A ferrule-capped optical fiber [MFC 400/430-0.66, 6-mm length, 400 μm-core, numerical aperture (NA) 0.48, MF 1.25 DoricLenses] was implanted 0.1 mm above the injection site and secured to the skull with black dental cement (Contemporary Ortho-JetLiquid, REF 1504-BL) and methyl methacrylate powder, Cooralite Dental 525000) and bone microscrews (00.90 × 0.063 M-SS-BS). For the chemogenetic inhibition 650 nl AAV5.hSyn.DIO.hM4D(Gi).mCherry or AAV-Flex-hM4D-Gi (*46*) was injected bilaterally into the MeA (AP: −1.58; L: ±1.89; DV: 5.47 mm). After stereotaxic surgeries, viruses were allowed to incubate for 2 weeks before the behavioral tests were performed.

2 weeks after stereotaxic injection of an inhibitory DREADDs (AAV-Flex-hM4D-Gi) or a control virus (AAV-Flex-mCherry) acute inhibition of the MeA^SST+^ neurons was performed before the exploratory behaviors. Thus, mice received an injection with either JHU 37160 (0.1-0.3 mg/Kg; 7198, TOCRIS) or saline (1% DMSO) 35 min prior to the EPM and OF tests and mice were evaluated for avoidance response.

### In vivo Fiber Photometry and Data Analysis

4 weeks after the beginning of the light protocol, and 2 weeks after stereotaxic surgery, fiber photometry recordings during behavior were performed. To reduce stress caused by the patch cord connection, mice were habituated for >1 week prior to behavior experiments. The day of the experiment, to reduce the stress caused by the patch cord connection, mice were first connected to a patch cord and placed in a new cage for 5 min before being introduced to the behavior arena. Then animals were introduced into the OF or the EPM arena for 5 min and fiber photometry recordings were performed as previously described (*47*). Briefly, GCaMP8m (i.e., calcium-dependent) and isosbestic (i.e., calcium-independent) excitation light was provided by 465- and 405-nm light-emitting diodes using the RZ10X processor (Tucker-Davis Technologies, Alachua, FL), and delivered to the animal via low-autofluorescence fiber optic patch cord and cannula (Doric Lenses, 400 μm-core; 0.66 NA), a fluorescence mini-cube, and a 1 × 1 commutator (Doric Lenses). Emission light was collected through the same optic fiber, detected, and amplified by an RZ10X processor (Tucker-Davis Technologies, Alachua, FL). The frequency of stimulation (465 nm, 210 Hz; 405 nm, 330 Hz) and the real-time processing of emission signals were controlled, demodulated and low-pass–filtered at 6 Hz by RZ10X and Synapse software (Tucker-Davis Technologies). Fiber photometry fluorescence signals were processed using a customized Python script adapted from the work of Martianova et al. (*48*). In brief, raw data from the 465 and the 405 channels were extracted and down sampled at 10Hz. To correct for signal decay resulting from photobleaching, baseline fluorescence for each channel was identified using an adaptive iteratively reweighted penalized least squares algorithm and then subtracted from each channel. Each signal was standardized by subtracting the median response and dividing by the standard deviation. To correct for motion artifacts or non–Ca^2+^-dependent responses, we used non-negative robust linear regression to fit the standardized 405-nm signal to the standardized 465-nm signal. The normalized ΔF/F was calculated by subtracting the fitted 405-nm signal from the standardized 465-nm signal. To correlate the photometry signals with behavior, behavioral experiments were recorded using a video camera, and the location and activity of the mice were automatically tracked using EthoVision XT (Noldus Information Technology, USA). Peri-event time plots were created using the TTL timestamps generated automatically using the Noldus USB-IO box (Noldus Information Technology, USA). Only zone entrance events with intervals longer than 3 s and 10 s were included in the open field and EPM analysis respectively, to avoid overlapping entrance events. The mean, maximum value, minimum value and area under the curve (AUC) for desired exploratory events were calculated using the mean, max, min and trapezoid function from the NumPy Python library (*49*). Additionally, the amplitude was calculated by subtracting the mean value to the maximum value registered during each event.

### Immunohistochemistry and RNAscope *in situ* hybridization

Mice were deeply anesthetized and transcardially perfused with 1% phosphate-buffered saline (PBS) solution immediately followed by 4% paraformaldehyde solution in PBS (pH = 7.4). Then, the brains were collected, post-fixed in 4% PFA for 24 h, cryoprotected in 30% sucrose solution, and stored at 4°C. Frozen brains were sectioned (30 μm) with a standard Leica microtome (SM2010R) and stored in cryoprotectant solution at −20°C until use. For immunohistochemistry (IHC) analysis, six coronal brain sections encompassing the MeA were processed, and two or three sections were used for RNAscope in situ hybridization (RNA-ISH). IHC was performed in free-floating slices that were first blocked for 1 h in 0.1 M phosphate buffer (PB) containing 3% normal horse serum (NHS) and then permeabilized in 0.2% Triton X-100. Slices were then incubated with the following antibodies (Table 1) in 1% PBS, 3% NHS, and 0.2% Triton X-100 for 24 or 48 h at 4°C with constant shaking. After three washes in PBS, slices were incubated for 1 h at room temperature with fluorescent-tagged secondary antibodies (Table 2). Brain sections were washed, mounted in gelatin on glass slides, counter-stained, and cover-slipped with Dapi Fluoromount-G™ (Electron Microscopy Sciences, United States). RNA-ISH [Multiplex Fluorescent Reagent Kit version 2, Advanced Cell Diagnostics (ACD)] was performed following the manufacturer’s instructions for the probes specified in table 3 of the supplementary material. Sections were washed in PBS, treated with hydrogen peroxide and mounted in glass slides with 1% PBS. Later, brain sections were incubated in the retrieval buffer and dehydrated in ethanol, previously to a protease treatment. Then, sections were incubated with the specific probes for 2 h at 40°C, and the amplification steps were followed as indicated by the manufacturers. Finally, sections were counter-stained, and cover-slipped with Dapi Fluoromount-G™ (Electron Microscopy Sciences, United States). Images were taken with a Leica TCS SP8 multiphoton confocal microscopy system (Leica Microsystems, Germany) and LASX software (Leica Microsystems, Germany). Quantifications were performed blind with Fiji ImageJ software. Neuronal quantifications were performed blind with ImageJ software. Quantification data were plotted either as the average of positive neurons per nucleus per animal. Semiquantitative RNA-ISH scoring methodology was used to quantify Kcnj3 mRNA expression. A score of 0 means no staining or less than 1 dot for every neuron, whereas a score of 4 means greater than 15 dots per neuron/cell or > 10% dots in clusters.

#### Brain Samples for Single-nuclei RNA sequencing

Brains were rapidly extracted, cooled in ice-cold DMEM medium and placed, ventral surface up, into a chilled stainless steel brain matrix (catalog no. SA-2165, Roboz Surgical Instrument Co., Gaithersburg, MD). Using GFP fluorescence to demarcate the MeA, brains were blocked to obtain a single coronal section containing the entire MeA, ∼2 mm thick. The MeA was microdissected by knife cuts at its visually approximated dorsolateral borders and pooled by experimental condition. We pooled tissue from 3 female and 3 male mice for each experimental group (Control, LCD-BL and LCD-RBL). Pooled tissues were then snap-frozen (at –30 °C).

### Single-cell library preparation, sequencing and alignment

snRNA-seq libraries from MeA tissues were generated by the Center for Epigenomics, UCSD. Frozen tissues were processed to isolate nuclei using dounce homogenization. Briefly, samples were suspended in 1 mL of douncing buffer (0.25M sucrose (S1888, Sigma), 25 mM KCl (AM9640G, Invitrogen), 5 mM MgCl_2_ (194698, Mp Biomedicals Inc), 10 mM Tris-HCl pH 7.5 (15567027, Thermo Fisher Scientific), 0.1% Triton-X-100 (Sigma-Aldrich, T8787), 1X cOmplete EDTA-free protease inhibitor (05056489001, Roche), 1 mM DTT, 0.5 U/µL RNase inhibitor (Promega, N2515), and molecular biology water (46000-CM, Corning)) and transferred to a pre-chilled dounce homogenizer. Homogenization was carried out on ice first using the loose pestle (5–10 strokes), followed by the tight pestle (15–25 strokes) until organoids were uniformly disrupted. The homogenate was filtered through a 30 µm filter and the dounce was rinsed with ∼500 µL of douncing buffer, which was added to the homogenate. The homogenate was centrifuged at 1000 × g for 10 min at 4°C (run speed 3/3). The supernatant was discarded, and the pellet was resuspended in 1 mL of douncing buffer. The homogenate was centrifuged again at 1000 x g for 10 min at 4°C (run speed 3/3). The supernatant was discarded and the homogenate was resuspended in 400 µL of Sort Buffer (1 mM EDTA (Invitrogen, 15575020), 1 U/µL RNase inhibitor (Promega, N2515), 1% BSA (Sigma-Aldrich, SRE0036) in PBS). The tube and filter were rinsed with an additional 400 µL of Sort Buffer, then the homogenate was stained with DRAQ7 (1:100; Cell Signaling, 7406) on ice for 10 min. 120,000 nuclei were sorted using a SH800 sorter (Sony) into 90 µL of 5X collection buffer (1 U/µL RNase inhibitor (Promega, N2515), 5% BSA (Sigma-Aldrich, SRE0036) in PBS); the FACS gating strategy sorted based on particle size and DRAQ7 fluorescence. Sorted nuclei were centrifuged at 500 rcf for 5 min (4°C, run speed 3/3) and supernatant was removed. The nuclei were resuspended in 15 µL of reaction buffer (1% BSA (Sigma-Aldrich, SRE0036) in PBS, 1 U/uL RNase inhibitor (Promega, N2515), stained with 0.4% Trypan Blue, and counted on a hemocytometer. A total of 12,000 nuclei per reaction were loaded onto a Chromium Controller (10x Genomics) using the Chromium Next GEM Single Cell 3ʹ GEM Kit v3.1 (10x Genomics, 1000123) and Chromium Next GEM Single Cell 3ʹ Gel Bead Kit v3.1 (10x Genomics, 1000122). Libraries were generated according to manufacturer recommendations using the Chromium Next GEM Single Cell 3ʹ Library Kit v3.1 (10x Genomics, 1000157), Chromium Next GEM Chip G Single Cell Kit (10x Genomics, 1000127), and Single Index Kit T Set A (10x Genomics, 1000213) for sample indexing. SPRISelect reagent (Beckman Coulter, B23319) was used for size selection and clean-up steps. Final library concentration was assessed by Qubit dsDNA HS Assay Kit (Thermo Fisher Scientific) and fragment size was checked using TapeStation High Sensitivity D1000 (Agilent) to ensure that fragment sizes were normally distributed about 500 bp. Libraries were sequenced using the NextSeq500 and a NovaSeq 6000 (Illumina) with the following read lengths: 28 + 8 + 91 (Read1 + Index1 + Read2).

### Alignment of snRNA-seq

snRNA libraries for each of the three conditions were aligned using the 10x Genomics CellRanger software (v.2.02, 10x Genomics) against the 10x Genomics pre-built mouse reference refdata-cellranger-mm10-2020-A-2.0.0 and quantified using “cellranger count”. Data from the three runs were aggregated using “cellranger-aggr” to normalize for sequencing depth.

### Quality control and preprocessing of snRNA-seq data

All snRNA-seq preprocessing was performed with CellRanger and Seurat v5. For each sample, we computed metrics for each cell including the number of unique genes detected (nFeature_RNA); the total molecules detected (nCount_RNA) and the percentage of reads mapping to the mitochondrial genome (percent.mt) (Suppl. Fig. 12A-C). We removed cells for which any of these metrics was more than three s.d. units from the mean in the sample. Next, we normalized the count data for each sample using SCTransform with percent.mt as a covariate. Principal component (PC) analysis was performed using the top 3,000 variable features with the top 50 PCs retained. The k-nearest neighbors (KNN) method was used to generate initial cell clusters, and a combination of SingleR and manual clustering based on established cell type markers were used to arrive at the final six primary cell type clusters. These results were used to calculate a uniform manifold approximation projection (UMAP) graph to represent the RNA and perform clustering. Subsetting of neurons and reclustering was performed to identify specific neuronal subtypes based on expression of *Gad2*, *Sst*, *Calb1*, *Slc17a6* and *Slc17a7*.

### Gene Ontology, Cell-cell communication, CellChat, and NeuronChat analysis

Gene ontology analysis was performed using the “enrichGO” package in R. Brain-specific cell-cell communication was analyzed using the NeuronChat analysis package in R (21). Cell clusters identified and annotated in Seurat were converted to NeuronChat objects using the workflow outlined in the NeuronChat vignettes. Briefly, the NeuronChat objects were created using the “createNeuronChat” function with data extracted from the “data” slot of the “RNA” assay from the associated Seurat object. The following additional parameters were specified: DB = “mouse”, assay = NULL, do.sparse = TRUE. Cell-cell communications were analyzed within each sample using the “run_NeuronChat” function with M=10. Visualizations of comparisons across groups were performed using the “net_aggregation” and “netVisual_circle_neuron” functions.

### Statistical analysis

Investigators who participated in end point analyses were blinded to the light protocol. Statistical analysis was carried out using GraphPad Prism (GraphPad Software, La Jolla, CA, United States).Outliers were identified and removed by the ROUT method provided by Graph Pad Prism Software, and normality was assessed (Shapiro–Wilk, Kolmogorov–Smirnov, D’Agostino and Pearson, and Anderson–Darling tests) before performing the corresponding statistical analyses. For normally distributed data, a parametric test was used one-way or two-way analysis of variance (ANOVA) or Student’s t-test and p-value less than 0.05 was considered significant, and was indicated as followed: *p < 0.05; **p < 0.01; ***p < 0.001; ****p < 0.0001. The statistical tests performed for each experiment are indicated in the figure legends.

## Supporting information

Supplementary Matherials

## Acknowledgements

We thank Korrus for providing the light prototype to perform the LCD-RBL experiment and Ben Harrison for setting up the prototype. We thank Sebastian Preissl from the Center for Epigenomics for technical assistance with the snRNA-seq library preparation. We thank Lily Quick and Jenna Brawley for their assistance with cell counting and all members of the Porcu’ Laboratory for valuable discussions and comments. This publication includes snRNA-sequencing data generated at the UCSD IGM Genomics Center which is partly funded by the UC San Diego School of Medicine.

## Author contributions

Conceptualization: A.P. Methodology: A.P., P.B., D.M., Investigation: P.B., D.M., A.S., J.K., A.B. Supervision: A.P. Writing (original draft): A.P. and P.B.

## Data and materials availability

All data needed to evaluate the conclusions in the paper are present in the paper and/or the Supplementary Materials. Raw data files from snRNAseq are uploaded to the Gene Expression Omnibus (GEO). Codes used for analysis in R and Python are available upon request.

## References

1. F. Falchi, P. Cinzano, D. Duriscoe, C. C. M. Kyba, C. D. Elvidge, K. Baugh, B. A. Portnov, N. A. Rybnikova, R. Furgoni, The new world atlas of artificial night sky brightness. Sci Adv 2, e1600377 (2016).

2. R. M. Lunn, D. E. Blask, A. N. Coogan, M. G. Figueiro, M. R. Gorman, J. E. Hall, J. Hansen, R. J. Nelson, S. Panda, M. H. Smolensky, R. G. Stevens, F. W. Turek, R. Vermeulen, T. Carreón, C. C. Caruso, C. C. Lawson, K. A. Thayer, M. J. Twery, A. D. Ewens, S. C. Garner, P. J. Schwingl, W. A. Boyd, Health consequences of electric lighting practices in the modern world: A report on the National Toxicology Program’s workshop on shift work at night, artificial light at night, and circadian disruption. Sci. Total Environ. 607-608, 1073–1084 (2017).

3. M. H. Smolensky, L. L. Sackett-Lundeen, F. Portaluppi, Nocturnal light pollution and underexposure to daytime sunlight: Complementary mechanisms of circadian disruption and related diseases. Chronobiol. Int. 32, 1029–1048 (2015).

4. T. A. Bedrosian, R. J. Nelson, Timing of light exposure affects mood and brain circuits. Transl. Psychiatry 7, e1017 (2017).

5. S. Tancredi, T. Urbano, M. Vinceti, T. Filippini, Artificial light at night and risk of mental disorders: A systematic review. Sci. Total Environ. 833, 155185 (2022).

6. M. Hysing, S. Pallesen, K. M. Stormark, R. Jakobsen, A. J. Lundervold, B. Sivertsen, Sleep and use of electronic devices in adolescence: results from a large population-based study. BMJ Open 5, e006748 (2015).

7. B. P. Hasler, R. E. Dahl, S. M. Holm, J. L. Jakubcak, N. D. Ryan, J. S. Silk, M. L. Phillips, E. E. Forbes, Weekend–weekday advances in sleep timing are associated with altered reward-related brain function in healthy adolescents. Biol. Psychol. 91, 334–341 (2012).

8. M. O. Mireku, M. M. Barker, J. Mutz, I. Dumontheil, M. S. C. Thomas, M. Röösli, P. Elliott, M. B. Toledano, Night-time screen-based media device use and adolescents’ sleep and health-related quality of life. Environ. Int. 124, 66–78 (2019).

9. A. L. Gamble, A. L. D’Rozario, D. J. Bartlett, S. Williams, Y. S. Bin, R. R. Grunstein, N. S. Marshall, Adolescent sleep patterns and night-time technology use: results of the Australian Broadcasting Corporation’s Big Sleep Survey. PLoS One 9, e111700 (2014).

10. D. Paksarian, K. E. Rudolph, E. K. Stapp, G. P. Dunster, J. He, D. Mennitt, S. Hattar, J. A. Casey, P. James, K. R. Merikangas, Association of Outdoor Artificial Light at Night With Mental Disorders and Sleep Patterns Among US Adolescents. JAMA Psychiatry 77, 1266–1275 (2020).

11. G. E. Guindon, C. A. Murphy, M. E. Milano, J. A. Seggio, Turn off that night light! Light-at-night as a stressor for adolescents. Front. Neurosci. 18, 1451219 (2024).

12. Y. M. Cissé, J. Peng, R. J. Nelson, Dim light at night prior to adolescence increases adult anxiety-like behaviors. Chronobiol. Int. 33, 1473–1480 (2016).

13. J. C. Borniger, Z. D. McHenry, B. A. Abi Salloum, R. J. Nelson, Exposure to dim light at night during early development increases adult anxiety-like responses. Physiol. Behav. 133, 99–106 (2014).

14. U. A. Tooley, D. S. Bassett, A. P. Mackey, Environmental influences on the pace of brain development. Nat. Rev. Neurosci. 22, 372–384 (2021).

15. A. C. Tonon, D. B. Constantino, G. R. Amando, A. C. Abreu, A. P. Francisco, M. A. B. de Oliveira, L. K. Pilz, N. B. Xavier, F. Rohrsetzer, L. Souza, J. Piccin, A. Caye, S. Petresco, P. H. Manfro, R. Pereira, T. Martini, B. A. Kohrt, H. L. Fisher, V. Mondelli, C. Kieling, M. P. L. Hidalgo, Sleep disturbances, circadian activity, and nocturnal light exposure characterize high risk for and current depression in adolescence. Sleep 45 (2022).

16. P. Bonilla, A. Shanks, Y. Nerella, A. Porcu, Effects of chronic light cycle disruption during adolescence on circadian clock, neuronal activity rhythms, and behavior in mice. Front. Neurosci. 18, 1418694 (2024).

17. S. M. Miller, D. Marcotulli, A. Shen, L. S. Zweifel, Divergent medial amygdala projections regulate approach–avoidance conflict behavior. Nat. Neurosci. 22, 565–575 (2019).

18. W. Hong, D.-W. Kim, D. J. Anderson, Antagonistic control of social versus repetitive self-grooming behaviors by separable amygdala neuronal subsets. Cell 158, 1348–1361 (2014).

19. S. Hattar, M. Kumar, A. Park, P. Tong, J. Tung, K.-W. Yau, D. M. Berson, Central projections of melanopsin-expressing retinal ganglion cells in the mouse. J. Comp. Neurol. 497, 326–349 (2006).

20. T. A. LeGates, C. M. Altimus, H. Wang, H.-K. Lee, S. Yang, H. Zhao, A. Kirkwood, E. T. Weber, S. Hattar, Aberrant light directly impairs mood and learning through melanopsin-expressing neurons. Nature 491, 594–598 (2012).

21. R. Chakraborty, M. J. Collins, H. Kricancic, D. Moderiano, B. Davis, D. Alonso-Caneiro, F. Yi, K. Baskaran, The intrinsically photosensitive retinal ganglion cell (ipRGC) mediated pupil response in young adult humans with refractive errors. J Optom 15, 112–121 (2022).

22. L. S. Mure, Intrinsically Photosensitive Retinal Ganglion Cells of the Human Retina. Front. Neurol. 12, 636330 (2021).

23. K. S. Scherf, J. M. Smyth, M. R. Delgado, The amygdala: an agent of change in adolescent neural networks. Horm. Behav. 64, 298–313 (2013).

24. G. Vandewalle, C. Schmidt, G. Albouy, V. Sterpenich, A. Darsaud, G. Rauchs, P.-Y. Berken, E. Balteau, C. Degueldre, A. Luxen, P. Maquet, D.-J. Dijk, Brain responses to Violet, blue, and green monochromatic light exposures in humans: Prominent role of blue light and the brainstem. PLoS One 2, e1247 (2007).

25. G. Vandewalle, S. Schwartz, D. Grandjean, C. Wuillaume, E. Balteau, C. Degueldre, M. Schabus, C. Phillips, A. Luxen, D. J. Dijk, P. Maquet, Spectral quality of light modulates emotional brain responses in humans. Proc. Natl. Acad. Sci. U. S. A. 107, 19549–19554 (2010).

26. V. Pilorz, S. K. E. Tam, S. Hughes, C. A. Pothecary, A. Jagannath, M. W. Hankins, D. M. Bannerman, S. L. Lightman, V. V. Vyazovskiy, P. M. Nolan, R. G. Foster, S. N. Peirson, Melanopsin Regulates Both Sleep-Promoting and Arousal-Promoting Responses to Light. PLoS Biol. 14, e1002482 (2016).

27. S. Jin, M. V. Plikus, Q. Nie, CellChat for systematic analysis of cell-cell communication from single-cell transcriptomics. Nat Protoc, doi: 10.1038/s41596-024-01045-4 (2024).

28. Y. E. Wu, L. Pan, Y. Zuo, X. Li, W. Hong, Detecting Activated Cell Populations Using Single-Cell RNA-Seq. Neuron 96, 313–329.e6 (2017).

29. W. Zhao, K. G. Johnston, H. Ren, X. Xu, Q. Nie, Inferring neuron-neuron communications from single-cell transcriptomics through NeuronChat, bioRxiv (2023). 10.1101/2023.01.12.523826.

30. B. N. Vo, E. Marron Fernandez de Velasco, T. R. Rose, H. Oberle, H. Luo, C. R. Hopkins, K. Wickman, Bidirectional influence of limbic GIRK channel activation on innate avoidance behavior. J. Neurosci. 41, 5809–5821 (2021).

31. S. J. Crowley, S. W. Cain, A. C. Burns, C. Acebo, M. A. Carskadon, Increased sensitivity of the circadian system to light in early/mid-puberty. J. Clin. Endocrinol. Metab. 100, 4067–4073 (2015).

32. Mechanisms mediating the effects of light on sleep and alertness: current challenges. Current Opinion in Physiology 15, 152–158 (2020).

33. M. A. Siraji, V. Kalavally, A. Schaefer, S. Haque, Effects of Daytime Electric Light Exposure on Human Alertness and Higher Cognitive Functions: A Systematic Review. Front. Psychol. 12, 765750 (2022).

34. G. Wang, Y.-F. Liu, Z. Yang, C.-X. Yu, Q. Tong, Y.-L. Tang, Y.-Q. Shao, L.-Q. Wang, X. Xu, H. Cao, Y.-Q. Zhang, Y.-M. Zhong, S.-J. Weng, X.-L. Yang, Short-term acute bright light exposure induces a prolonged anxiogenic effect in mice via a retinal ipRGC-CeA circuit. Sci Adv 9, eadf4651 (2023).

35. O. Lawal, F. P. Ulloa Severino, C. Eroglu, The role of astrocyte structural plasticity in regulating neural circuit function and behavior. Glia 70, 1467–1483 (2022).

36. Y. Ota, A. T. Zanetti, R. M. Hallock, The role of astrocytes in the regulation of synaptic plasticity and memory formation. Neural Plast. 2013, 185463 (2013).

37. N. A. Perez-Catalan, C. Q. Doe, S. D. Ackerman, The role of astrocyte-mediated plasticity in neural circuit development and function. Neural Dev. 16, 1 (2021).

38. S. Paixão, R. Klein, Neuron-astrocyte communication and synaptic plasticity. Curr. Opin. Neurobiol. 20, 466–473 (2010).

39. C. Lüscher, P. A. Slesinger, Emerging roles for G protein-gated inwardly rectifying potassium (GIRK) channels in health and disease. Nat. Rev. Neurosci. 11, 301– 315 (2010).

40. A. Albrecht, M. Thiere, J. R. Bergado-Acosta, J. Poranzke, B. Müller, O. Stork, Circadian modulation of anxiety: a role for somatostatin in the amygdala. PLoS One 8, e84668 (2013).

41. K. Yu, P. Garcia da Silva, D. F. Albeanu, B. Li, Central Amygdala Somatostatin Neurons Gate Passive and Active Defensive Behaviors. J. Neurosci. 36, 6488–6496 (2016).

42. J. M. Stujenske, P.-K. O’Neill, C. Fernandes-Henriques, I. Nahmoud, S. R. Goldburg, A. Singh, L. Diaz, M. Labkovich, W. Hardin, S. S. Bolkan, T. R. Reardon, T. J. Spellman, C. D. Salzman, J. A. Gordon, C. Liston, E. Likhtik, Prelimbic cortex drives discrimination of non-aversion via amygdala somatostatin interneurons. Neuron 110, 2258– 2267.e11 (2022).

43. Y. Sun, L. Qian, L. Xu, S. Hunt, P. Sah, Somatostatin neurons in the central amygdala mediate anxiety by disinhibition of the central sublenticular extended amygdala. Mol. Psychiatry, doi: 10.1038/s41380-020-00894-1 (2020).

44. S. Ahrens, M. V. Wu, A. Furlan, G.-R. Hwang, R. Paik, H. Li, M. A. Penzo, J. Tollkuhn, B. Li, A Central Extended Amygdala Circuit That Modulates Anxiety. J Neurosci 38, 5567–5583 (2018).

45. R. K. Butler, L. C. White, D. Frederick-Duus, K. F. Kaigler, J. R. Fadel, M. A. Wilson, Comparison of the activation of somatostatin- and neuropeptide Y-containing neuronal populations of the rat amygdala following two different anxiogenic stressors. Exp. Neurol. 238, 52–63 (2012).

46. B. L. Roth, DREADDs for Neuroscientists. Neuron 89, 683–694 (2016).

47. A. Porcu, A. Nilsson, S. Booreddy, S. A. Barnes, D. K. Welsh, D. Dulcis, Seasonal changes in day length induce multisynaptic neurotransmitter switching to regulate hypothalamic network activity and behavior. Sci Adv 8, eabn9867 (2022).

48. E. Martianova, S. Aronson, C. D. Proulx, Multi-Fiber Photometry to Record Neural Activity in Freely-Moving Animals. J. Vis. Exp., doi: 10.3791/60278 (2019).

49. C. R. Harris, K. J. Millman, S. J. van der Walt, R. Gommers, P. Virtanen, D. Cournapeau, E. Wieser, J. Taylor, S. Berg, N. J. Smith, R. Kern, M. Picus, S. Hoyer, M. H. van Kerkwijk, M. Brett, A. Haldane, J. F. Del Río, M. Wiebe, P. Peterson, P. Gérard-Marchant, K. Sheppard, T. Reddy, W. Weckesser, H. Abbasi, C. Gohlke, T. E. Oliphant, Array programming with NumPy. Nature 585, 357–362 (2020).

