## Supplementary Matherials for "Evening blue light exposure during adolescence induces avoidance behaviors and rewires medial amygdala circuit"

**Supplementary materials**

**Supplementary Figure 1. Blue light and reduced blue light spectrum and chromaticity.** The light source was characterized for its spectral properties and chromaticity using a spectroradiometer. The spectral distribution (top-right panel) shows a peak wavelength (λp) and a peak intensity (λpV). The correlated color temperature (CCT) was measured at 6322 K for **A** and 4057K for **B**, with a Duv value of 0.0179 for **A** and 0.0377 for **B**. The chromaticity coordinates were plotted on the CIE 1931 (bottom left) and CIE 1976 (bottom right) diagrams. XYZ tristimulus values (X = 11966.07, Y = 14368.00, Z = 3021.44) provide additional information on the light's intensity and colorimetric properties. These data describe the spectral and chromatic properties of the light sources used in the LCD protocols.

**Supplementary Figure 2. LCD-RBL Show No Effect on Circadian Locomotor Activity and activity in a new environment. (A)** Representative actograms showing wheel running activity under LCD-RBL conditions for 3 weeks on adolescent mice. The yellow background represents when the light is on, the green background represents when the reduced blue light (RBL) is on, and the dark background represents the dark phase. (**B)** Total activity counts in LCD-RBL mice during the weekdays and weekends. (**C)** Onset of the LCD-RBL mice during the weekdays and weekends [$F_{(1, 24)} = 5.156$, $p < 0.0324$ by two-way ANOVA with Šídák's multiple comparison posttest]. (**D)** Total activity counts of the LCD-RBL mice during the weekdays and weekends. Individual data points represent independent mice, data are shown as mean ± SEM. (**E)** Distance traveled in the Open Field (OF) test by control, LCD-BL and LCD-RBL mice. Individual data points represent independent mice, data are shown as mean ± SEM. One-way ANOVA Tukey multiple comparison post-test.

**Supplementary Figure 3. LCD-BL exposure does not increase avoidance behaviors in adult mice.** Bar graphs show time spent **i**n the inner zone (**A**) and in the periphery (**B)** of the open field arena. ns: Not significant. Bar graphs show time spent in the open arms **C)** and in the closed arms (**D)** of the elevated plus maze. Individual data points represent independent mice, data are shown as mean ± SEM. Student's t-test. ns: Not significant.

**Supplementary Figure 4.** Feature plots showing expression of literature reported marker genes for cell clusters in the snRNA-seq data.

**Supplementary Figure 5. (A)** Representative confocal micrographs showing the neuronal marker NeuN (red) detected by immunofluorescence in the MeA of adolescent mice (Scale bar 100 µm). (**B)** Bar graph shows the number of NeuN positive cells in control, LCD-BL, and LCD-RBL. (**C**) Representative confocal micrographs showing the astrocytic marker glial fibrillary acidic protein (GFAP) (red) detected by immunofluorescence in the MeA of adolescent mice (Scale bar 100 µm). (**D)** Bar graph shows number of GFAP positive cells in control, LCD-BL, and LCD-RBL.

Individual data points represent independent mice, data are shown as mean ± SEM. One-way ANOVA Tukey multiple comparison post-test: *$p < 0.05$.

**Supplementary Figure 6.** Heatmap showing the expression levels of selected top cell-type marker genes (fold change > 3, FDR < 0.05) across individual cells.

**Supplementary Figure 7. GO terms of molecular functions in neuronal clusters (A**) Top gene ontology terms (FDR < 0.15) upregulated and downregulated in the neuronal population in LCD-BL vs control. (**B**) Top gene ontology terms upregulated and downregulated in the neuronal population in LCD-RBL vs LCD-BL.

**Supplementary Figure 8. LCD-BL reduces Kcnj3 expression in the MeA$^{SST+}$ neurons.** Representative confocal micrographs showing *Kcnj3* (red) mRNA expression detected by RNAscope in (**A)** SST neurons (*SST*, blue), (**C)** GABAergic neurons (*VGAT*, green) and (**E)** glutamatergic neurons (*VGLUT2*, blue) in the MeA (Scale bar 20 µm). Bar graphs show *Kcnj3* mRNA expression determined by semiquantitative scoring of *Kcnj3* dots and clusters per (**B**) SST neuron, (**D)** GABAergic neurons and **(E)** glutamatergic neurons in the MeA of adolescent mice exposed to control, LCD-BL and LCD-RBL conditions. Individual data points represent independent mice, data are shown as mean ± SEM. One-way ANOVA Tukey multiple comparison post-test: *$p < 0.05$, **$p < 0.01$, ***$p < 0.001$.

**Supplementary Figure 9.** Violin plots show GCaMP8m (**A)** Maximum value recorded, (**B)** Minimum value recorded and (**C)** Amplitude before (Pre-Exploration) and **(D)** Maximum value recorded, **(E)** Minimum value recorded and (**F)** Amplitude and during the exploration (Exploration) in the open field test. Violin plots show **(G)** Maximum value recorded, **(H)** Minimum value recorded and (**I)** Amplitude before (Pre-Exploration) and (**J)** Maximum value recorded, **(K)** Minimum value recorded and (**L)** Amplitude and during the exploration (Exploration) in the elevated plus maze test. Individual data points represent independent mice, data are shown as mean ± SEM. Student's t-test. ns: Not significant.

**Supplementary Figure 10.** Distance travelled in the open field test (**A)** and elevated plus maze test (**B)** after JHU or saline injection. Individual data points represent independent mice, data are shown as mean ± SEM. One-way ANOVA Tukey multiple comparison post-test: **$p < 0.01$

**Supplementary Figure 11. (A)** Representative confocal micrographs showing the cFOS (green) detected by immunofluorescence in the MeA of adolescent mice (Scale bar 100 µm). **(B)** Bar graph shows c-FOS expression in the MeA of adolescent mice after open field and elevated plus maze tests. Individual data points represent independent mice, data are shown as mean ± SEM. One-way ANOVA Tukey multiple comparison post-test: **$p < 0.01$, ***$p < 0.001$.

**Supplementary Figure 12.** snRNA-seq profiles of 6 mice per experimental group. For each experimental group the number of unique genes detected per cell (nFeature_RNA) (**A**), total number of reads within each cell (nCount_RNA) (**B**), and percentage of percent mitochondrial reads (**C**) are shown for each cell. Quality metrics calculated with Seurat v5.

**Table 1. Primary antibodies**

| Source | Primary antibody | Concentration | REF |
|---|---|---|---|
| Chicken | NeuN | 1:500 | Millipore, ABN81 |
| Chicken | GFAP | 1:500 | Invitrogen, PA1-10004 |
| Chicken | GFP | 1:500 | Invitrogen, A10262 |
| Rabbit | Somatostatin | 1:1000 | Invitrogen, PA5-85759 |
| Rabbit | c-Fos | 1:500 | Cell Signaling, 2250S |
| Guinea Pig | c-Fos | 1:1000 | Synaptic Systems, 226308 |

**Table 2. Secondary antibodies**

| Source | Primary antibody | Concentration | REF |
|---|---|---|---|
| 647 nm | anti-rabbit | 1:1000 | Invitrogen, A31573 |
| 555 nm | anti-chicken | 1:1000 | Invitrogen, A21437 |
| 488 nm | anti-chicken | 1:1000 | Invitrogen, A78948 |
| 594 nm | anti -GuineaPig | 1:1000 | Jackson Immuno, 706-585-148 |
| gp | c-Fos | 1:1000 | Synaptic Systems, 226308 |
| 647 nm | anti-rabbit | 1:1000 | Invitrogen, A31573 |

**Table 3. RNAscope probes**

| Probe | Concentration | REF |
|---|---|---|
| c-FOS | | 316921 |
| Kcnj3 | | 523951 |
| VGLUT2 | 1:30 | 319171-C2 |
| SST | 1:100 | 404631-C2 |
| VGAT | 1:40 | 319191-C3 |
